# COMPARATIVE MINERAL COMPOSITION ANALYSIS OF COMMON (ROASTED) AND GREEN (UNROASTED) BUCKWHEAT (*FAGOPYRUM ESCULENTUM*)

**DOI:** 10.1101/2025.10.09.681340

**Authors:** Hamzayeva Nargiza Rajabboyevna, Ishimov Uchkun Jomurodovich

## Abstract

This study comparatively evaluated the elemental composition of green (raw) and common (thermally processed) *Fagopyrum esculentum* using inductively coupled plasma mass spectrometry (ICP-MS). Finely ground samples of both types were acid-digested and analyzed for macroelements (Ca, Mg, Na, K), microelements (Fe, Zn, Cu, Mn, Co, Ni, Cr, Mo), trace elements (Se, La, Zr, Y), and heavy metals (Ba, Cd, As, Sb). Results revealed that green buckwheat contained significantly higher levels of macroelements, particularly Mg (3130 ppm), Ca (5061 ppm), and Na (1500 ppm), compared to common buckwheat. Iron content was notably elevated in green buckwheat (1234 ppm) relative to the processed form (22 ppm), with moderate increases in Mn, Cr, and Zn. Trace elements such as La, Ce, and Y were also more abundant, while all heavy metals remained within WHO/FAO safety limits. These findings indicate that green buckwheat retains a richer and more diverse mineral profile due to minimal processing and may serve as a nutritionally superior source of essential elements for functional food development and micronutrient deficiency prevention.

## Introduction

Buckwheat (*Fagopyrum esculentum*) is a pseudocereal that has garnered considerable scientific interest in recent years due to its high nutritional value, absence of gluten, and a wide range of bioactive compounds associated with various health-promoting effects. In contrast to conventional cereals, buckwheat (*Fagopyrum esculentum*) is distinguished by its high content of essential amino acids, antioxidants, dietary fiber, and a diverse array of bioactive compounds, including rutin, quercetin, and various phenolic acids. These phytochemicals are associated with significant therapeutic potential, particularly in the prevention and management of cardiovascular diseases, diabetes mellitus, obesity, and inflammatory conditions (Sofi et al., 2022), (Kaur et al., 2023).

An important component of the nutritional evaluation of buckwheat is its mineral profile, encompassing both macroelements (e.g., sodium, magnesium, potassium, calcium) and microelements (e.g., iron, zinc, copper, manganese). These minerals perform essential physiological functions, including participation in enzymatic reactions, oxygen transport, bone mineralization, and the regulation of metabolic processes (Alghamdi et al., 2023). In addition, rare trace elements such as selenium (Se), vanadium (V), and molybdenum (Mo), though required in trace amounts, are indispensable for maintaining antioxidant defense mechanisms and modulating immune responses. Conversely, the presence of toxic heavy metals, including lead (Pb) and cadmium (Cd), even at low concentrations, poses significant health hazards and necessitates stringent monitoring, particularly in food crops cultivated on contaminated soils (Tchounwou et al., 2012). The concentrations of toxic heavy metals, including cadmium (Cd), arsenic (As), and barium (Ba), detected in both common and green buckwheat samples were found to be well below the maximum permissible limits established by international food safety authorities such as the World Health Organization (WHO) and the Food and Agriculture Organization (FAO) of the United Nations. This indicates that the levels present pose no significant health risk and confirms that both types of buckwheat are safe for regular human consumption from a toxicological standpoint (Bawwab et al., 2022). Beyond its micronutrient composition, buckwheat has garnered attention for its potential effects on satiety—the physiological sensation of fullness following food intake. Its high dietary fiber content and rich mineral profile are hypothesized to modulate satiety-related hormones and appetite-regulating pathways, thereby positioning buckwheat as a promising dietary component in the management of body weight and the prevention of metabolic syndrome (Begum et al., 2025).

Despite the widespread consumption of buckwheat, comparative investigations into the elemental composition and toxicological profiles of common (roasted) versus green (raw) buckwheat remain scarce. Such studies are essential to comprehensively evaluate their respective nutritional benefits and potential health risks (Bonafaccia et al., 2003).

This study aims to address the existing gap in the literature by evaluating and comparing the concentrations of essential macroelements (such as sodium, potassium, calcium, and magnesium), microelements (including iron, zinc, copper, and manganese), as well as trace elements in both common (roasted) and green (unroasted) buckwheat. In addition to compositional analysis, the study explores the physiological implications of these elements, particularly their roles in modulating satiety, appetite regulation, and metabolic health.

## Materials and Methods

### Sample Preparation

Samples of common (thermally processed) and green (raw) buckwheat (*Fagopyrum esculentum*) were obtained from certified local suppliers. All samples were air-dried at room temperature, finely ground using a laboratory mill, and stored in airtight containers until analysis. Prior to elemental analysis, the samples were homogenized and passed through a 1 mm sieve to ensure uniformity.

As shown in Figure 1 the grains exhibit a characteristic dark brown coloration due to thermal processing,which may alter their chemical composition, including the content and activity of bioactive compounds such as flavonoids and antioxidants.

**Figure 1.**
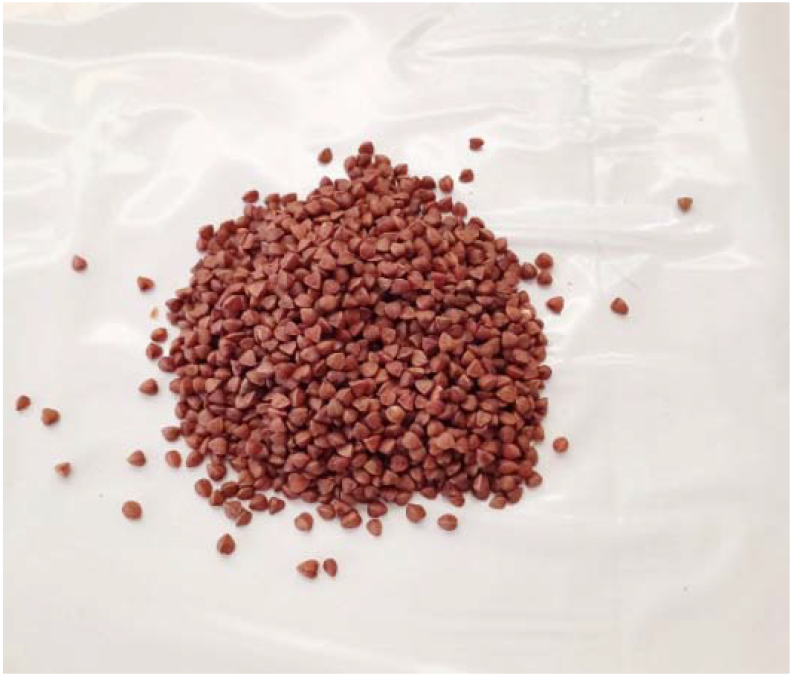
Roasted buckwheat *(Fagopyrum esculentum)* grains

Figure 2 illustrates light-colored grains have not undergone thermal treatment and thus retain their native biochemical profile, making them suitable for studies focused on the extraction and characterization of thermolabile bioactive compounds.

**Figure 2.**
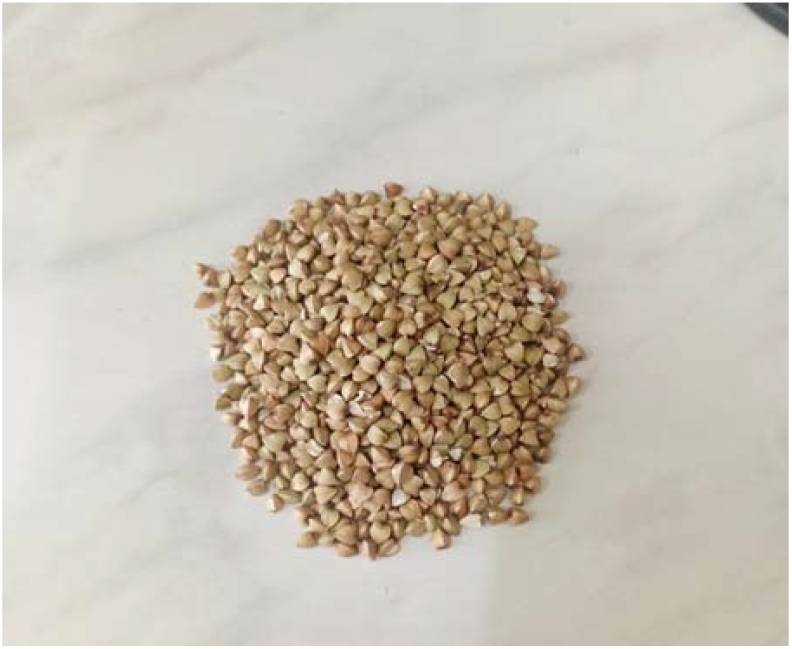
Green (unroasted) buckwheat *(Fagopyrum esculentum)* grains

### Elemental Analysis via Inductively Coupled Plasma Mass Spectrometry (ICP-MS)

The determination of macroelements (Ca, Mg, Na, K), microelements (Fe, Zn, Cu, Mn, Co, Ni, Cr, Mo), trace elements (Ce, La, Zr, Mo, Y), and heavy metals (Sb, Cd, As) was carried out using inductively coupled plasma mass spectrometry (ICP-MS). For sample digestion, approximately 0.0500 to 0.5000 g of each ground sample was accurately weighed using an analytical balance (±0.0001 g) and placed into teflon digestion vessels.

A mixture of concentrated nitric acid (HNO□, ≥69%, suprapur grade) and hydrogen peroxide (H□O□, 30%, analytical grade) was added to each vessel. The samples were then digested using a Berghof MWS-3+ microwave digestion system (Germany), following a program optimized for organic matrices. After complete digestion, the clear solutions were cooled, transferred quantitatively into 50 or 100 mL volumetric flasks, and diluted to volume with 0.5% (v/v) nitric acid.

Elemental concentrations were measured using a NexION 2000 ICP-MS system (PerkinElmer, USA), operated with high-purity argon gas (≥99.995%) as the plasma source. The instrument was calibrated using a certified multi-element standard solution (Standard No. 3), containing 29 elements relevant for food analysis. Bidistilled deionized water was used for all dilutions and blank preparations. Internal standards were applied to correct for matrix effects and instrumental drift. All reagents and materials were of analytical or trace metal grade quality. Quality control was ensured through the use of method blanks and certified reference materials (CRMs).

## Results and discussion

The inductively coupled plasma mass spectrometry (ICP-MS) analysis revealed significant differences in the elemental composition between common and green buckwheat samples. Green buckwheat demonstrated notably higher concentrations of most macro-, micro-, heavy metal, and trace elements compared to common buckwheat.

According to Figure 3 illustration, among macroelements, green buckwheat showed elevated levels of magnesium (Mg: 3130 ppm), calcium (Ca: 5061 ppm), and sodium (Na: 1500 ppm), which were substantially higher than in common buckwheat (Mg: 2543 ppm, Ca: 2869 ppm, Na: 989 ppm). Potassium (K) levels also exhibited moderate increases.

**Figure 3.**
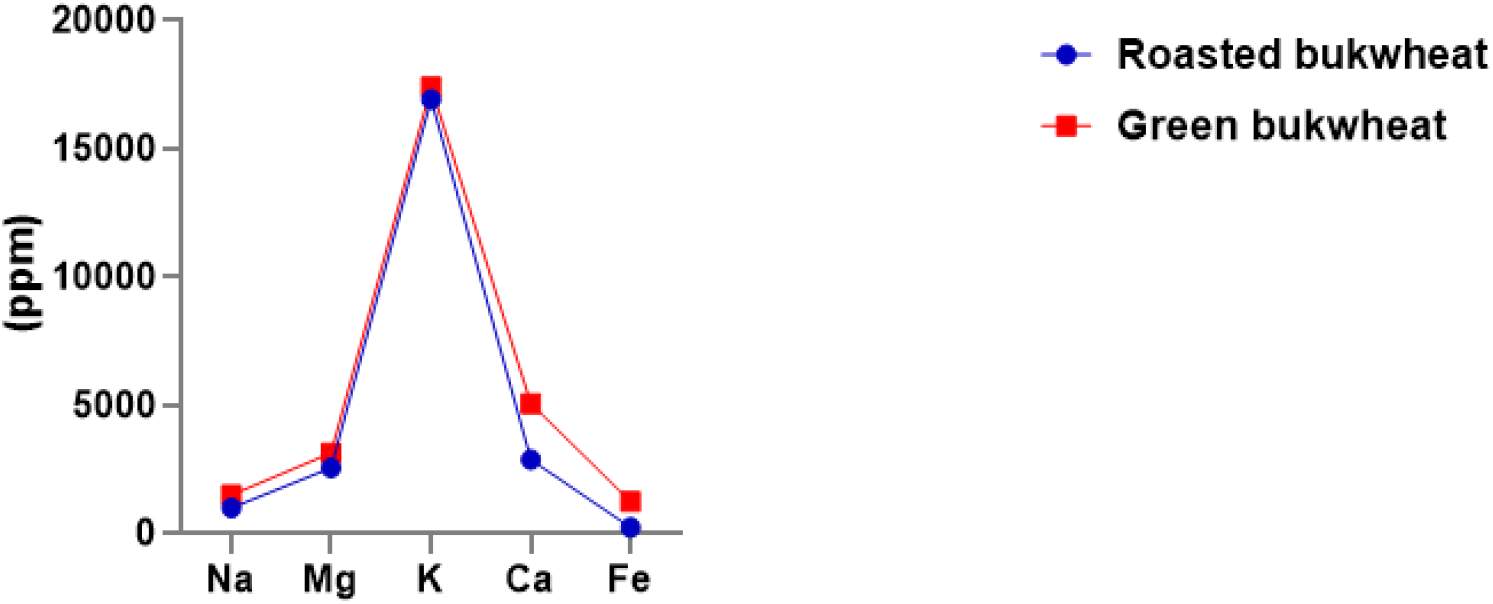
The comparison of mineral composition between common roasted buckwheat and green buckwheat shows significant differences in macroelement concentration

As shown in Figure 4 regarding microelements, iron (Fe) concentration in green buckwheat reached 1234 ppm, which is dramatically higher than in common buckwheat (22 ppm), indicating its potential as a superior dietary iron source. Zinc (Zn), manganese (Mn), and chromium (Cr) were also present in significantly greater amounts in green buckwheat (Zn: 47.3 ppm, Mn: 174.3 ppm, Cr: 4.45 ppm) compared to common buckwheat (Zn: 48.3 ppm, Mn: 145.4 ppm, Cr: 3 ppm). The analysis of toxic heavy metals showed that barium (Ba) was higher in green buckwheat (7.33 ppm) than in common buckwheat (4.83 ppm). However, cadmium (Cd), arsenic (As), and antimony (Sb) levels were below toxic thresholds in both samples, with Sb remaining below the detection limit (<0.30 ppm).

**Figure 4.**
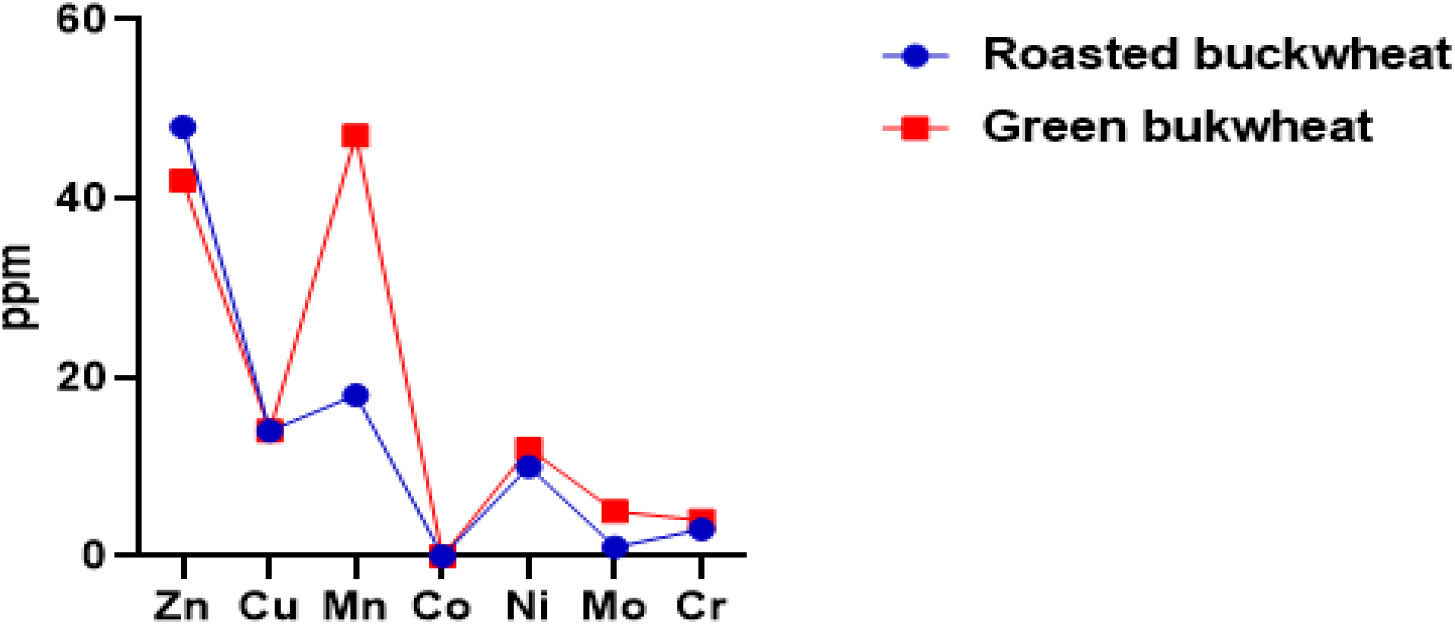
The bar chart illustrates the concentrations of various microelements in common (roasted) buckwheat and green (unroasted) buckwheat

In the Table 1 demonstration, trace elements were also more abundant in green buckwheat. Notably, lanthanum (La), cerium (Ce), and yttrium (Y) concentrations were substantially higher (La: 0.265 ppm, Ce: 0.596 ppm, Y: 0.371 ppm) than in common buckwheat, where most values were at or below detection limits. These findings indicate that green buckwheat has a richer and more diverse mineral profile, suggesting its enhanced nutritional and functional food potential compared to common buckwheat.

**Table 1.** The table presents the concentration of toxic heavy metals and trace elements in both common and green buckwheat.

| Toxic heavy metal elements |  |  | Trace elements |  |  |
| --- | --- | --- | --- | --- | --- |
| Element name | Common buckwheat | Green buckwheat | Element name | Common buckwheat | Green buckwheat |
| Ba | 4.83 | 7.33 | La | 0.04 | 0.265 |
| Cd | 0.054 | 0.04 | Ce | 0.084 | 0.596 |
| As | 0.139 | 0.189 | Y | <0,10 | 0,371 |
| Sb | <0,30 | <0,30 | Zr | <0,10 | <0,10 |

The present study provides a comprehensive comparison of the mineral composition of common and green buckwheat using ICP-MS. The findings demonstrate that green buckwheat contains significantly higher concentrations of essential macroelements such as magnesium (Mg), calcium (Ca), potassium (K), and sodium (Na), which are vital for numerous physiological functions including bone health, nerve transmission, and electrolyte balance. Notably, Mg and Ca were especially abundant in green buckwheat, highlighting its potential as a mineral-dense functional food.

Among microelements, green buckwheat showed a remarkable enrichment in iron (Fe), with levels exceeding 50-fold compared to common buckwheat. This suggests that green buckwheat may serve as a valuable dietary source of iron, which is essential for oxygen transport and cellular metabolism, especially in populations vulnerable to iron deficiency anemia. The higher content of zinc (Zn), manganese (Mn), and chromium (Cr) in green buckwheat also indicates potential benefits for immune support, antioxidant defense, and glucose metabolism. The presence of trace elements such as lanthanum (La), cerium (Ce), and yttrium (Y) in green buckwheat, though typically found in very small amounts, may reflect its unique uptake capacity from the soil or differences in seed processing. These elements, while not considered essential, are increasingly being investigated for their biological roles and therapeutic potential in trace amounts.

Furthermore, the detection of trace levels of elements such as lanthanum (La), cerium (Ce), and neodymium (Nd) in green buckwheat adds a novel aspect to its nutritional and functional potential. Although present in minute concentrations, these elements have been associated with modulating oxidative stress pathways and enhancing neuroprotective functions in some recent studies (Hao et al., 2024). Their presence could offer added value to green buckwheat, potentially positioning it as a functional food with trace element-based bioactivity. Moreover, the higher concentrations of essential minerals like magnesium, potassium, and iron in green buckwheat compared to roasted samples further underscore the detrimental effect of thermal processing on mineral integrity. These elements play critical roles in cardiovascular health, glucose metabolism, and immune regulation (Bo & Pisu, 2008), (Szklarz et al., 2022). Therefore, promoting the consumption of minimally processed buckwheat may provide more consistent mineral bioavailability and contribute to improved public health nutrition.

Although both buckwheat types were within acceptable safety thresholds for toxic heavy metals (As, Cd, Sb), as determined by international regulatory standards (e.g., WHO/FAO), green buckwheat demonstrated a cleaner profile. This highlights its potential not only as a safe dietary source but also as a candidate for organic and export-quality food production, particularly in regions emphasizing clean-label and environmentally friendly agriculture. Importantly, levels of toxic heavy metals such as cadmium (Cd), arsenic (As), and antimony (Sb) remained below safety thresholds in both varieties, suggesting that neither sample poses a significant health risk from contamination. However, slightly higher concentrations of barium (Ba) and arsenic in green buckwheat highlight the need for continued monitoring, especially when buckwheat is cultivated in mineral-rich or polluted soils. The observed differences in mineral composition could be attributed to genetic variation, differences in seed coat permeability, or differential nutrient mobilization during germination and growth. Green buckwheat, often less processed and closer to its raw form, may retain more nutrients compared to thermally treated or roasted common buckwheat.

### Role of Buckwheat in Satiety

Buckwheat demonstrated a notable effect on short-term satiety, which may be attributed to its high content of resistant starch, dietary fiber, and bioactive flavonoids known to influence appetite regulation (Heidi Stirling, Stephanie Keddy, Jennifer Jamieson, Shannan Grant, Bohdan L Luhovyy, n.d.).

Buckwheat crackers did not lead to significant changes in acute blood glucose or insulin levels, they effectively influence the release of gastrointestinal hormones associated with satiety, indicating their potential in appetite regulation mechanisms(Stringer et al., 2013). Population-based studies have demonstrated an inverse relationship between dietary fiber consumption and body weight, with enhanced satiety and reduced energy intake proposed as potential mediating factors (Clark, M. J., & Slavin, J. L., 2013).

Overall, the superior mineral profile of green buckwheat underscores its value as a nutrient-dense food ingredient with potential applications in dietary interventions, especially in populations with mineral deficiencies. Further studies, including bioavailability assessments and long-term nutritional impacts, are needed to validate these findings and support its wider use in health-promoting food formulations.

## Conclusion

This study provides a comprehensive comparison of the mineral composition of common (roasted) and green (unroasted) buckwheat *(Fagopyrum esculentum)* using inductively coupled plasma mass spectrometry (ICP-MS). The results demonstrate that green buckwheat retains significantly higher levels of essential macro- and microelements, including magnesium, potassium, iron, and zinc, compared to roasted samples. The thermal processing involved in roasting appears to reduce the overall mineral content, likely due to volatilization, structural breakdown, or formation of less bioavailable complexes. Notably, the presence of trace rare earth elements in green buckwheat introduces a novel nutritional dimension, with potential implications for oxidative stress modulation and neuroprotection, although further biological studies are required to substantiate these effects. Both buckwheat types were found to be within safe limits for toxic elements such as arsenic, cadmium, and antimony, confirming their suitability for human consumption.

In summary, green buckwheat presents superior nutritional value from a mineralogical perspective and may offer enhanced functional food potential. The findings support the recommendation to promote the use of minimally processed buckwheat in health-conscious diets and emphasize the need for further research on mineral bioavailability and physiological impact in vivo.

